# From Inhibition to Excitation and Why: The Role of Temporal Urgency in Modulating Corticospinal Activity

**DOI:** 10.1101/2023.12.27.573452

**Authors:** Aaron N. McInnes, Benjamin Smithers, Ottmar V. Lipp, James R. Tresilian, Ann-Maree Vallence, John C. Rothwell, Welber Marinovic

**Author notes:** Corresponding Authors: Aaron N. McInnes, Welber Marinovic.

## Abstract

Previous research on movement preparation identified a period of corticospinal suppression about 200 ms prior to movement initiation. This phenomenon has been observed for different types of motor tasks typically used to investigate movement preparation (e.g., reaction time, self-initiated, and anticipatory actions). However, we recently discovered that this phenomenon is not observed when actions must be initiated under time pressure. In the present study, we investigated urgency effects on corticospinal suppression throughout the time course of an anticipatory timing task. Participants were required to perform timing actions under two urgency scenarios, high and low, and we applied single-pulse transcranial magnetic stimulation at different times during the time course of preparation. We analysed the time course of excitability under high and low scenarios in relation to expected and actual movement onset times. Our results confirmed our earlier findings that corticospinal suppression is not observed when participants perform actions under high urgency scenarios. In addition, we found no evidence that this preparatory suppression could be shifted in time to occur later under high urgency scenarios. Moreover, we found evidence that responses prepared under high urgency are more likely to be disrupted by external events (e.g., TMS pulses). These results suggest that preparatory suppression might be a strategy employed by the central nervous system to shield motor actions from interference of external events (e.g., loud sounds) when time allows. Given these data, we propose conceptual models that could account for the absence of preparatory suppression under time pressure to act.

## 1.0 Introduction

Although the initiation of motor actions seems simple to us in most instances, the central nervous system undergoes a series of intricate neural processes before an action unfolds. The dynamic of these processes occurring before movement initiation is reflected in changes in corticospinal (CS) excitability that can be measured using transcranial magnetic stimulation (TMS). A well-documented phenomenon when humans engage in action preparation is preparatory suppression. Preparatory suppression reflects a period of CS suppression relative to baseline (at rest) that is observed approximately 250 ms to 100 ms prior to action initiation (Davranche et al., 2007; Duque et al., 2010; Duque & Ivry, 2009; Hannah et al., 2018; Hasbroucq et al., 1997, 1999; Ibáñez et al., 2020; Klein et al., 2016; Lebon et al., 2016; Marinovic et al., 2015; McInnes et al., 2021; Starr et al., 1988; Zaaroor et al., 2003). Previous research investigating the time course of this preparatory suppression in reactive, anticipatory, and self-timed actions, showed that this period of CS suppression is followed by a period of facilitation, and this was common to all three types of movements participants performed (Ibáñez et al., 2020).

While previous studies indicate that CS suppression might be a necessary component of action preparation (Ibáñez et al., 2020), we recently reported data that contradicts this view (McInnes et al., 2021). We demonstrated that when temporal constraints on action preparation are high (e.g., short preparation times), the CS system is not suppressed during preparation. In more detail, when participants were given only 350 ms of movement preparation (high-urgency), there was no observable suppression of motor evoked potentials (MEPs) by single-pulse TMS 250 ms prior to movement onset. In contrast, the typical MEP suppression was observed at the same time-point when participants were given 1400 ms to prepare their action (low-urgency).

These findings not only suggest that CS suppression may be an unnecessary component of action preparation but, by observing the resulting effects on motor performance (e.g., peak force and rate of force development), also shed light on the potential function of premovement CS suppression. These data indicated that the effects of TMS on the execution of the prepared motor action depended on whether the system underwent preparatory CS suppression. During high-urgency preparation, when no preparatory CS suppression was observed, TMS impaired motor performance. In contrast, when CS suppression was observed, TMS facilitated motor performance of the prepared action as evidenced by an increase in response force. We also observed this differential effect of urgency on motor performance when a loud acoustic stimulus was presented instead of TMS pulses. Therefore, we proposed that preparatory CS suppression may protect prepared motor actions from interference by external stimuli.

While we observed no evidence of CS suppression 250 ms prior to action initiation during high-urgency preparation, we could not rule out that CS suppression is adaptively shifted to occur closer to the movement’s onset when preparation time is constrained. For example, neural signatures recorded from non-human primates indicate that action preparation can take place at a significantly shorter time-scale when actions must be prepared and initiated rapidly (Lara et al., 2018). Furthermore, when forced to prepare and initiate movements on a short time-scale, humans can move accurately in shorter durations of time than their typical reaction times would suggest (Haith et al., 2016). As such, it is possible that while CS suppression cannot be detected 250 ms prior to movement under high temporal constraints, there may be a rapid inhibition-disinhibition process that occurs within the 250 ms prior to action initiation when actions are prepared with high urgency.

Here, we probed CS excitability at two time points before movement initiation under high and low temporal constraints to examine the potential existence of a rapid inhibition-disinhibition process. We predicted that, in line with our previous report, there would be a CS inhibition at 250 ms prior to action initiation under low temporal constraints but not under high temporal constraints. Furthermore, we predicted that under both high and low temporal constraints, there would be a facilitation of CS excitability relative to baseline when probed 150 ms prior to action initiation. Such an observation would support the notion that CS suppression is not a necessary component of action preparation. To assess the relationship between urgency, CS suppression, and motor output, we also quantified motor output as a function of CS excitability and urgency. This quantification of preparatory parameters is beneficial in forming a theoretical understanding of the process of action preparation in healthy individuals and may be informative for the understanding of motor impairment due to neurological conditions. In addition, we also developed a simple model where excitatory and inhibitory processes develop during preparation, interacting to yield a net excitatory state which depends on preparation urgency.

## 2.0 Method

### 2.1 Participants

Twenty-two participants were recruited (15 female, mean age = 22.7, SD = 6.13, range = 18 – 46). Participants were screened in accordance with the TMS safety guidelines by Rossi et al. (2021) to identify potential contraindications. They were self-reportedly right-handed with no apparent neurological conditions or injuries which may have impacted their performance in the task. The experimental protocol was approved by Curtin University’s human research ethics committee and all participants provided informed, written consent in accordance with the declaration of Helsinki.

### 2.2 Procedures

TMS was administered during an anticipatory timing task under high and low temporal constraints, a protocol of which we have described previously (McInnes et al. 2021). Before beginning experimental trials, participants completed 10 practice trials (five trials for each urgency condition), we located the scalp location of the first dorsal interosseous (FDI) hotspot, and identified the resting motor threshold via the Rossini-Rothwell procedure (Rossini et al., 1994). This procedure involved applying single-pulse TMS over the FDI hotspot to identify the minimum intensity at which the TMS elicits an MEP of at least 50 µV in the target muscle in at least 5 out of 10 consecutive trials. During the experiment, participants were presented with the anticipatory task visual stimuli on a 24.5 inch monitor (Asus ROG PG258Q; 240 Hz refresh rate, 1920 × 1080 resolution) via custom MATLAB (Mathworks, v2015b) scripts written using Psychtoolbox (Kleiner et al., 2007, v3.0.11). Participants were presented with a clock on-screen and in each trial, after a delay period of a duration randomised from a uniform distribution of 3 – 5 s, the clock would begin revolving. Participants were instructed to initiate a ballistic abduction of the FDI in synchrony with the time at which the clock completed a full 360° revolution. The speed of the clock revolution was adjusted in each block so that a full revolution would take either 350 ms (high-urgency condition) or 1400 ms (low-urgency condition). Participants completed a block of 120 trials for each urgency condition, the order of which was counterbalanced between participants. In each block, there were four TMS conditions (TMS_BASELINE_, TMS_-250_, TMS_-150_, or control). In the TMS_BASELINE_ condition (20 trials), single TMS pulses were triggered while the participant was at rest, two seconds after the beginning of the delay period that preceded the start of the anticipatory clock’s revolution. In the TMS_-250_ condition (20 trials), the TMS pulse was triggered 250 ms prior to the time at which the anticipatory clock completed a full revolution (the expected time of movement onset). In the TMS_-150_ condition, the TMS pulse was triggered 150 ms prior to the expected time of movement onset (20 trials). In the control condition (60 trials), no TMS pulses were triggered. Trials were pseudo-randomized so that TMS was not presented in any two consecutive trials.

### 2.3 Transcranial magnetic stimulation

A Magstim D70^2^ figure-of-eight coil, with a wing diameter of 70 mm and connected to a Magstim BiStim^2^ stimulator (Magstim, Whitland, UK) was used to deliver TMS. The intensity of the stimulus was set for each participant at 120% of their individual resting motor threshold (to the nearest 1% of maximum stimulator output). The mean resting motor threshold was 39.75% of maximum stimulator output (range = 25% – 55%). TMS was triggered during the experiment from MATLAB via the MAGIC toolbox (v0.2, Habibollahi Saatlou et al., 2018).

### 2.4 Data acquisition, processing, and analysis

#### 2.4.1 Acquisition of electromyogram and force data

Electromyogram (EMG) and force data were digitised over the course of each trial at a sampling rate of 2 kHz with a National Instruments USB-6229 data acquisition device (National Instruments, Austin TX, USA). EMG data were recorded from the right FDI using bipolar 8 mm Ag/AgCl electrodes placed along the muscle belly and with a reference electrode placed on the styloid process of the right ulna. These data were amplified with a gain of x1000, notch filtered at mains frequency (50 Hz) and band-pass filtered between 20 and 500 Hz (Digitimer NeuroLog, Letchworth Garden City, UK).

Force data were recorded from a capacitive force sensor (SingleTact 10N linearly calibrated sensor; Pressure Profile Systems, UK) which was embedded in a custom 3D-printed housing. Participants rested the lateral side of their index finger on this sensor for the duration of the experiment so that they pressed against the sensor when performing the ballistic FDI abduction.

#### 2.4.2 Processing of electromyogram data

Data were processed and analysed using R statistics (R Core Team, 2016). During processing, MEP peak-to-peak amplitudes were measured for each trial in which TMS was administered using an automated algorithm which we have described previously (McInnes et al., 2021). All trials were also visually inspected and peak-to-peak MEP amplitudes were manually marked if they had been mislabelled by the automated algorithm, or rejected if visual inspection of the EMG recording indicated that significant noise, artifacts, or voluntary activation of FDI may have impacted detection of peak-to-peak MEP amplitudes. The manual rejection of MEPs resulted in the removal of 449 trials in total (16.27% of all trials, participant median = 19, range = 1 - 47). In addition, after manual rejection of MEPs, the median root mean square (RMS) of EMG activity 200 ms prior to TMS presentation was calculated for each participant as a measure of baseline EMG activity. Excessive EMG activity during this baseline period may indicate the presence of slight voluntary activation of the FDI which may have impacted the size of the resulting MEP. Therefore, MEPs were rejected if their baseline EMG RMS exceeded this median RMS value by three standard deviations. This resulted in the rejection of an additional 46 MEPs (1.99% of trials, participant median = 0, range = 0 - 15). After filtering, 2265 MEPs remained for analysis.

#### 2.4.3 Processing of force data

Force recordings from each trial were detrended and multiplied by a factor of 10 to convert the voltage output of the capacitive force sensor to Newtons (N). These data were subject to Teasdale’s (1993) algorithm to detect the time of movement onset. The expected time of movement onset (the time at which the anticipatory clock completed a full revolution) was subtracted from the observed time of movement onset for each trial to determine the temporal error of movement onset. Peak force was determined as the peak of the force signal following movement onset. Control trials were excluded from analyses if their temporal error of movement onset relative to the expected time of movement onset was < -150 ms or > 150 ms, or if no movement onset could be detected. Trials for which TMS was presented were not filtered based on this criterion so that we could examine TMS effects on action initiation in contrast to appropriate task performance in the absence of TMS (control trials). The rationale behind this criterion is to guarantee that we only analysed control trials in which the temporal accuracy of the response was within an acceptable level of performance (e.g., participants were not too late or early due to inattention). This resulted in the removal of 297 trials (4.97% of all behavioural trials). After filtering, 5675 behavioural trials remained for analysis.

#### 2.4.4 Data analyses

We conducted a series of linear mixed-effects models to examine the effect of TMS presentation on motor output (temporal error of movement onset, voluntary response vigour, and voluntary peak response force) under preparation urgency conditions (high urgency, low urgency) and with subject IDs as a random effect. To examine the effects of preparation urgency on MEP amplitudes over the course of preparation, a generalised linear mixed-effects model was run with subject IDs as a random effect, MEP amplitude as a response variable, and with preparation urgency (low, high), and TMS timing (TMS_BASELINE_, TMS_-250_, TMS_-150_) as predictor variables. Statistical significance was determined at *a* = .05 and *R*^2^ values are reported as an estimate of effect size. We also calculated the amplitude of each TMS_-250_ and TMS_-150_ trial MEP as a percentage of the median amplitude of TMS_BASELINE_ MEPs, for each urgency condition. These data were analysed using a generalised linear mixed-effects model with subject IDs as a random effect, % TMS_BASELINE_ MEP amplitude as an outcome variable, and with preparation urgency and TMS timing (TMS_-250_, TMS_-150_) as dependent variables.

To examine changes in CS excitability during preparation relative to action initiation, we conducted additional analyses with data realigned so that the time of TMS presentation was relative to the observed time of movement onset (rather than the expected time of movement onset/cue). Trials for which TMS was presented < -350 ms or > 0 ms relative to the onset of the voluntary action were discarded for these analyses. To this end, we analysed MEP amplitudes as a percentage of baseline and peak rate of force development, with the time of TMS presentation relative to movement onset and foreperiod duration as predictor variables, and subject IDs as a random effect. Given the non-linearity of these outcome measures over time, we conducted these analyses using generalized additive models, with cubic spline smoothing over time and across foreperiod durations to examine differences in CS excitability over the course of preparation under different preparatory temporal constraints.

## 3.0 Results

### 3.1 Temporal error of movement onset

Analysis of the temporal error of movement onset across foreperiod durations and TMS timings indicated a significant main effect of TMS timing, *F*_(3, 5679.4)_ = 10.8, *p* < .001, *R*^2^ = .006. Temporal error of movement onset was earlier in TMS_BASELINE_ (M = -41.15 ms, SD = 296.78, *p* < .001), TMS_-250_ (M = -38.97 ms, SD = 156.7, *p* < .001), and TMS_-150_ (M = -31.86 ms, SD = 145.25, *p* = .024) trials in comparison to the Control condition (M = -14.96 ms, SD = 50.51). Temporal error did not differ between the TMS_-250_ and TMS_-150_ conditions (*p* = .691), the TMS_-250_ and TMS_BASELINE_ conditions (*p* = .997), or the TMS_-150_ and TMS_BASELINE_ conditions (*p* = .553). In the linear model of temporal error, the main effect of foreperiod duration, *F*_(1, 5679.3)_ = 0.33, *p* = .563, *R*^2^ = .000, and the interaction of foreperiod duration with TMS timing, *F*_(3, 5678.9)_ = 0.63, *p* = .593, *R*^2^ = .000, were not significant. A Bayesian t-test of subject mean temporal error in control trials between the two foreperiod durations provided substantial evidence (Jeffreys, 2006) against an effect (BF_01_ = 6.18) of foreperiod duration on temporal error. Temporal error of movement onset is plotted in **FIGURE 1**.

**FIGURE 1.**
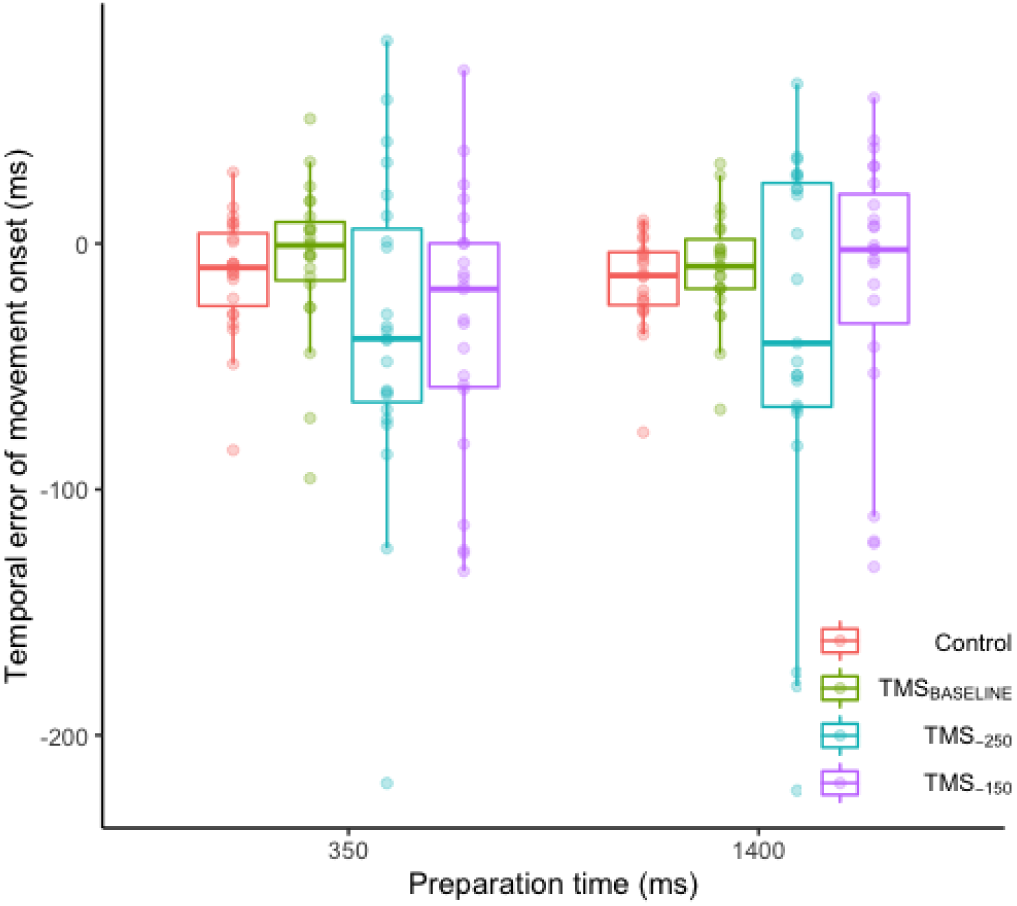
Temporal error of movement onset across foreperiod durations and for trials in each TMS timing condition. Boxplots depict trial-level median and interquartile range, individual points represent subject means.

### 3.2 Stimulation timing differentially affects motor output as a function of urgency

We examined output of voluntary actions (peak force and peak rate of force development) as a function of TMS condition and foreperiod duration. The main effects of TMS, *F*_(3, 5679)_ = 25.7, *p* < .001, *R*^2^ = .013, and foreperiod duration, *F*_(1, 5679)_ = 56.57, *p* < .001, *R*^2^ = .01, each had a significant effect on log-normalised peak force. Log-normalised peak force was greatest on average in control trials (*M* = 0.84, *SD* = 0.77), and was significantly greater than responses in TMS_-250_ trials (*M* = 0.65, *SD* = 0.80, *p* < .001), and TMS_-150_ trials (*M* = 0.77, *SD* = 0.77, *p* = .003), but not TMS_BASELINE_ trials (*M* = 0.81, *SD* = 0.75, *p* = .694). Peak force was reduced from TMS_BASELINE_ in TMS_-250_ trials (*p* < .001) but not TMS_-150_ trials (*p* = .224). The difference was also significant between TMS_-250_ and TMS_-150_ trials (*p* < .001). For the effect of foreperiod duration on log-normalised peak force, motor output was greater, on average, in the high urgency condition (*M* = 0.87, *SD* = 0.73) in comparison to the low urgency condition (*M* = 0.72, *SD* = 0.8, *p* < .001). However, the interaction of foreperiod duration with TMS condition was not significant on log-normalised peak force *F*_(3, 5679)_ = 1.81, *p* = .143, *R*^2^ = .000. Peak force as a function of TMS condition and foreperiod duration is plotted in **FIGURE 2A**. Ratios of peak force in TMS trials relative to control trials indicated a significant main effect of TMS condition, *F*_(2, 2633.6)_ = 3.67, *p* = .026, *R*^2^ = .003, with peak force ratios differing between TMS_BASELINE_ (*M* = 1.12, *SD* = 2.76) and TMS_-250_ (*M* = 0.76, *SD* = 2.81, *p* = .024) conditions, but not between TMS_BASELINE_ and TMS_-150_ conditions (*M* = 0.86, *SD* = 3.36, *p* = .143) or between TMS_-250_ and TMS_-150_ conditions (*p* = .734). Peak force ratios as a function of TMS condition and foreperiod duration are plotted in **FIGURE 2B**. The main effect of foreperiod duration, *F*_(1, 2633.3)_ = 0.96, *p* = .327, *R*^2^ = .000, and the interaction of TMS condition with foreperiod duration, *F*_(2, 2633.2)_ = 0.49, *p* = .616, *R*^2^ = .000, were not significant.

**Figure 2.**
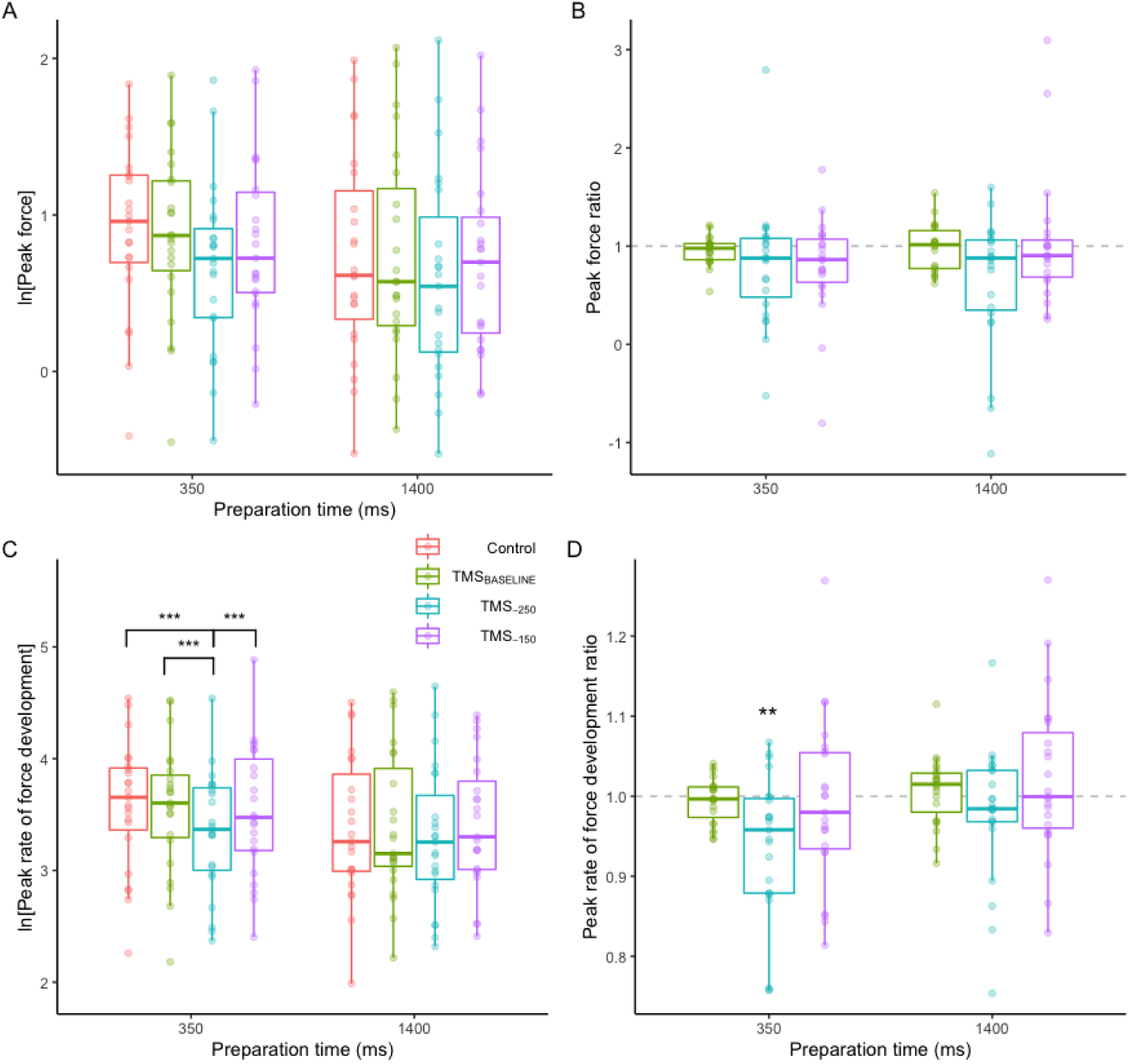
Motor output as a function of the time of TMS delivery and preparation urgency. A).Log-normalised peak force across TMS conditions and foreperiod durations. B). Peak force ratios (TMS trials relative to control) across TMS conditions and foreperiod durations. C).Log-normalised peak rate of force development across TMS conditions and foreperiod durations. D). Peak rate of force development ratios (TMS trials relative to control) across TMS conditions and foreperiod durations, with significance depicted against a mean of 1. Box plots represent median and interquartile range. Individual points represent individual subject averages. *** = *p* < .001, ** = *p* < .01, *p* < .05.

For the model of log-normalised peak rate of force development, the main effects of TMS condition *F*_(3, 5679)_ = 20.77, *p* < .001, *R*^2^ = .011, and foreperiod duration, *F*_(1, 5679)_ = 67.92, *p* < .001, *R*^2^ = .012, were both significant. In addition, the interaction of TMS condition with foreperiod duration was significant in the model, *F*_(3, 5679)_ = 5.22, *p* = .001, *R*^2^ = .003. Post-hoc tests of the main effects of TMS condition indicated peak rate of force development was reduced in the TMS_-250_ condition (*M* = 3.33, *SD* = 0.76), relative to control trials (*M* = 3.5, *SD* = 0.75, *p* < .001), TMS_BASELINE_ trials (*M* = 3.49, *SD* = 0.76, *p* < .001), and TMS_-150_ trials (*M* = 3.48, *SD* = 0.77, *p* < .001). The difference in peak rate of force development was not significant between TMS_-150_ and control trials (*p* = .861), TMS_-150_ and TMS_BASELINE_ (*p* = .980), or TMS_BASELINE_ and control trials (*p* = .990). For the main effect of foreperiod duration on log-normalised peak rate of force development, response vigour was greater in the high urgency condition (*M* = 3.55, *SD* = 0.74) in comparison to the low urgency condition (*M* = 3.39, *SD* = 0.77, *p* < .001). Finally, post hoc tests of the interaction of TMS condition with foreperiod duration indicated TMS timing had differentially impacted response vigour, depending on preparation urgency. Log-normalised peak rate of force development was lower in TMS_-250_ trials (*M*_HighUrgency_ = 3.34, *SD*_HighUrgency_ = 0.76, *M*_LowUrgency_ = 3.32, *SD*_LowUrgency_ = 0.76) relative to control (*M*_HighUrgency_ = 3.6, *SD*_HighUrgency_ = 0.72, *p* < .001; *M*LowUrgency = 3.4, *SD*LowUrgency = 0.78, *p* = .094), TMSBASELINE (*M*HighUrgency = 3.58, *SD*_HighUrgency_ = 0.7, *p* < .001; *M*_LowUrgency_ = 3.4, *SD*_LowUrgency_ = 0.76, *p* = .205), and TMS_-150_ (*M*HighUrgency = 3.55, *SD*HighUrgency = 0.79, *p* < .001; *M*LowUrgency = 3.42, *SD*LowUrgency = 0.75, *p* = .068) trials for the high urgency condition, but not in the low urgency condition. Response vigour did not change in TMS_-150_ trials, relative to control trials, for either the high urgency (*p* = .534) or the low urgency conditions (*p* = .992). Similarly, response vigour was no different in TMS_-150_ trials, relative to TMS_BASELINE_, in either the high urgency (*p* = .973), or low urgency (*p* = .999) conditions. Finally, response vigour was not different in TMS_BASELINE_ trials, relative to control trials, in either the high urgency (*p* = .997), or low urgency (*p* = .999) conditions. Peak rate of force development as a function of TMS condition and foreperiod duration is plotted in **FIGURE 2C**. In addition, the model of peak rate of force development ratios for TMS trials, relative to control trials and across foreperiod durations indicated significant main effects of TMS condition *F*_(2, 2632.3)_ = 21.61, *p* < .001, *R*^2^ = .016, foreperiod duration *F*_(1, 2632.2)_ = 24.11, *p* < .001, *R*^2^ = .009, and an interaction of the two *F*_(2, 2632.2)_ = 3.4, *p* = .034, *R*^2^ = .003. Given that a ratio equal to 1 would indicate no change from control trials, as a follow-up test, we ran one-sample t-tests against a mean of 1 for peak rate of force development ratios across TMS conditions and foreperiod durations, to examine evidence for facilitation or suppression of response vigour relative to control trials. For the high urgency condition, peak rate of force development ratios was significantly lower in the TMS_-250_ condition (*p* = .002), but not the TMS_BASELINE_ (*p* = .349) or TMS_-150_ (*p* = .638) conditions. In the low urgency condition, peak rate of force development ratios were not significantly different from a mean of 1 in any of the TMS_BASELINE_ (*p* = .476), TMS_-250_ (*p* = .238), or TMS_-150_ (*p* = .385) conditions. Peak rate of force development as a function of TMS condition and foreperiod duration is plotted in **FIGURE 2D**.

### 3.3 No evidence for suppression during high-urgency preparation

#### 3.3.1 Background EMG prior to TMS

A generalised linear mixed-effects model of EMG RMS error 200 ms prior to TMS presentation indicated no significant interaction of foreperiod duration with TMS timing, χ^2^_(2)_ = 0.95, *p* = .623, *R*^2^ = 0.010. However, the main effect of foreperiod duration was significant, χ^2^_(1)_ = 50.75, *p* < .001, *R*^2^ = 0.009, with RMS error being significantly greater for trials with the 350 ms foreperiod duration (2.20 × 10^-3^ mV) in comparison to the 1400 ms foreperiod duration (2.22 × 10^-3^ mV; *p* < .001). Mean RMS error is plotted in **FIGURE 3A**.

**FIGURE 3.**
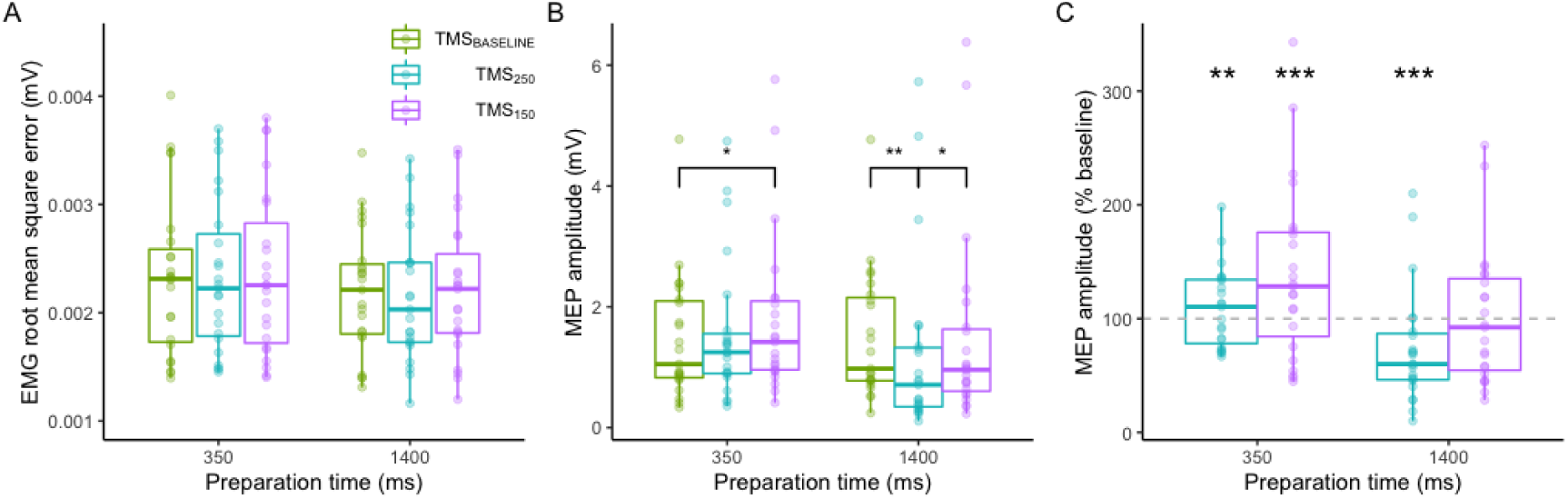
Cue-locked MEP amplitude. (A) EMG root mean square error 200 ms prior to TMS presentation across foreperiod durations and for each TMS timing condition. (B) MEP amplitude across foreperiod durations and each TMS timing condition. (C) MEP amplitude as a percentage of baseline across foreperiod durations and each TMS timing condition, significance indicates difference relative to baseline (100%). Dashed line represents 100% (no change from baseline). Boxplots depict median and interquartile range, individual points represent subject means. * = *p* < .05. ** = *p* < .01. *** = *p* < .001.

#### 3.3.2 Cue-locked motor evoked potentials

A generalised linear model of the amplitude of MEPs evoked by TMS showed a two-way interaction between foreperiod duration and TMS timing, χ^2^_(2)_ = 13.03, *p* = .001, *R*^2^ = 0.004. To account for the potential confound of differences in EMG RMS error prior to TMS presentation between foreperiod durations, we ran additional models which included EMG RMS error as a random effect. The interaction of foreperiod duration with TMS timing remained significant when including EMG RMS error as a random intercept and as a random slope. To compare these models, we employed the Akaike Information Criterion (AIC), a metric used to evaluate the quality of a model in relation to other potential models. The AIC of the model which contained only a random intercept for subject IDs was lower (AIC = 4848.1) compared to that of the models which also included a random EMG RMS error intercept (AIC = 4850.1) and slope (AIC = 4854.1). Therefore, we report results of the simpler model which included a random intercept for subject IDs only. Post-hoc tests indicated that MEP amplitudes in the TMS_BASELINE_ condition were not significantly different between the 1400 ms (*M* = 1.51 mV, *SD* = 1.32) and 350 ms (*M* = 1.46 mV, *SD* = 1.41) foreperiod durations (*p* = .734). However, for the 1400 ms foreperiod duration, MEP amplitudes were significantly lower in the TMS_-250_ condition (*M* = 1.14 mV, *SD* = 1.61; *p* = .003), but not the TMS_-150_ condition (*M* = 1.26 mV, *SD* = 1.62; *p* = .435), relative to the TMS_BASELINE_ condition (see **FIGURE 3**). MEP amplitudes were also significantly larger in the TMS_-150_ condition relative to the TMS_-250_ condition (*p* = .033). In contrast, for the 350 ms foreperiod duration, MEP amplitudes were no different from TMS_BASELINE_ in the TMS-_250_ condition (*M* = 1.55 mV, *SD* = 1.65; *p* = .285). Furthermore, TMS-_150_ MEP amplitudes (*M* = 1.71 mV, *SD* = 1.65) were significantly increased from TMS_BASELINE_ (*p* = .034). There was no significant difference between the TMS-_250_ and TMS-_150_ MEP amplitudes for the 350 ms foreperiod duration (*p* = .285). Mean MEP amplitudes are plotted in **FIGURE 3B**.

Analysis of the MEP amplitudes as a percentage of baseline indicated a significant interaction of TMS timing and foreperiod duration, χ^2^_(1)_ = 6.38, *p* = .012, *R*^2^ = 0.03. All pairwise comparisons were significant in the model. In addition, we conducted a t-test against a mean of 100 to assess the evidence for suppression/facilitation of MEP amplitude percentages relative to baseline, for each TMS timing and foreperiod duration (with correction for the familywise error rate via the Holm procedure). For the 1400 ms foreperiod duration, MEP amplitudes as a percentage of baseline were significantly lower than a mean of 100% in the TMS_-250_ condition (*M* = 69.81%, *SD* = 70.92; *p* < .001), but not the TMS_-150_ condition (*M* = 94.42%, *SD* = 109.13; *p* = .340). For the 350 ms foreperiod duration, MEP amplitudes as a percentage of baseline were significantly greater than a mean of 100% in both the TMS_-250_ (*M* = 110.89, *SD* = 73.02; *p* = .008) and TMS_-150_ (*M* = 135.77, *SD* = 125.89; *p* < .001) conditions. MEP amplitudes as a percentage of baseline are plotted in **FIGURE 3C**.

#### 3.3.3 Response-locked motor evoked potentials

To examine changes in CS excitability relative to action initiation, we modelled MEP amplitudes with data realigned so that the time of TMS presentation was relative to the onset of the voluntary action. TMS was presented on average 211.69 ms prior to movement onset in TMS_-250_ trials, and 140.17 ms prior to movement onset in the TMS_-150_ condition. A generalised additive model of MEP amplitudes over time indicated a significant effect of foreperiod duration, χ^2^_(1)_ = 14.22, *p* < .001. Similarly, a generalised additive model of MEP amplitudes as a percentage of baseline over time indicated a significant main effect of foreperiod duration, χ^2^ = 43.75, *p* < .001. Modelled MEP amplitudes, MEP amplitudes as a percentage of baseline, and the distribution of trials across the time-series prior to movement onset are shown in **FIGURE 4A-C**.

**FIGURE 4.**
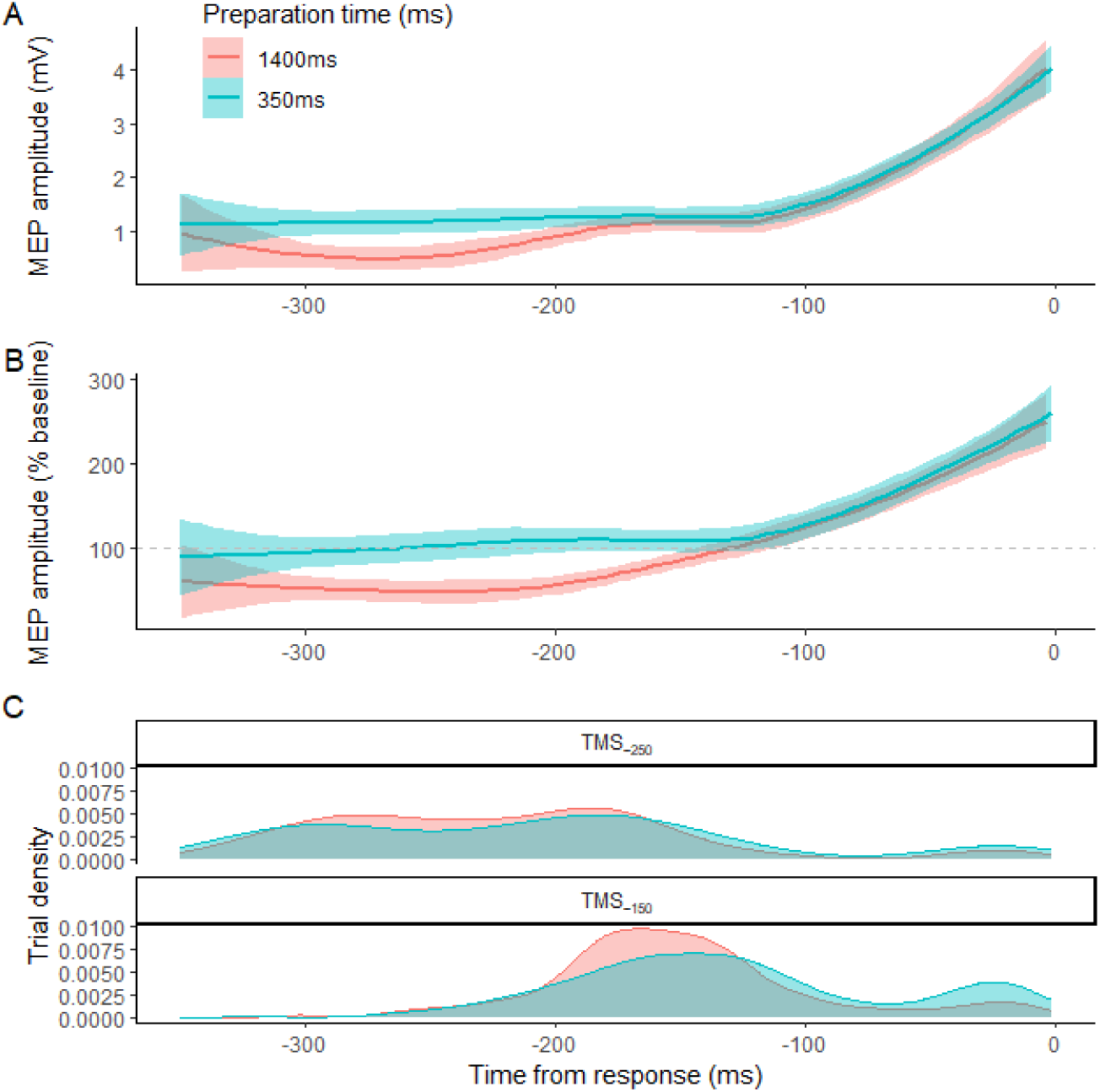
Response-locked MEP amplitude. (A) MEP amplitudes over time, relative to the onset of the voluntary action. (B) MEP amplitudes as a percentage of baseline over time, relative to the onset of the voluntary action. (C) Distribution of trial density as a function of the time of TMS presentation relative to onset of the voluntary action, across foreperiod durations and TMS timing conditions.

In addition, we modelled MEP amplitudes over time in the 1400 ms foreperiod duration condition using the generalised additive method, to estimate non-linear changes in CS excitability over the course of preparation under temporal constraints typically observed in the laboratory. MEP amplitudes predicted using this model against observed values had a *R*^2^ = .37 and Pearson’s *r* = .61. The derivative of these modelled data indicated that there were two significant periods of change, the first occurring approximately 238 – 182 ms prior to movement onset, and the second occurring approximately 118 – 4 (end of the time-series) ms prior to movement onset.

### 3.4 Motor output is modulated as a function of time of TMS presentation and urgency

We examined motor output with TMS presentation relative to movement onset as a continuous predictor. The main effect of TMS presentation relative to movement onset on peak rate of force development ratios was not significant, χ^2^_(1)_ = 0.33, *p* = .566. However, the interaction of TMS presentation relative to movement onset with foreperiod duration was significant, χ^2^_(1)_ = 3.87, *p* = .049, supporting TMS’ effects on motor output over the course of preparation depending on preparation urgency. The correlation between peak rate of force development ratios and TMS timing relative to movement onset was significant and positive for the low urgency preparation condition, *r* = .08, *p* = .019, suggesting that motor output was facilitated in trials where TMS was presented closer to movement onset under low urgency preparation. However, the correlation was not significant for the high urgency preparation condition, *r* = -.03, *p* = .448. Peak rate of force development ratios as a function of TMS presentation relative to movement onset and across foreperiod durations is plotted in **FIGURE 5**.

**Figure 5.**
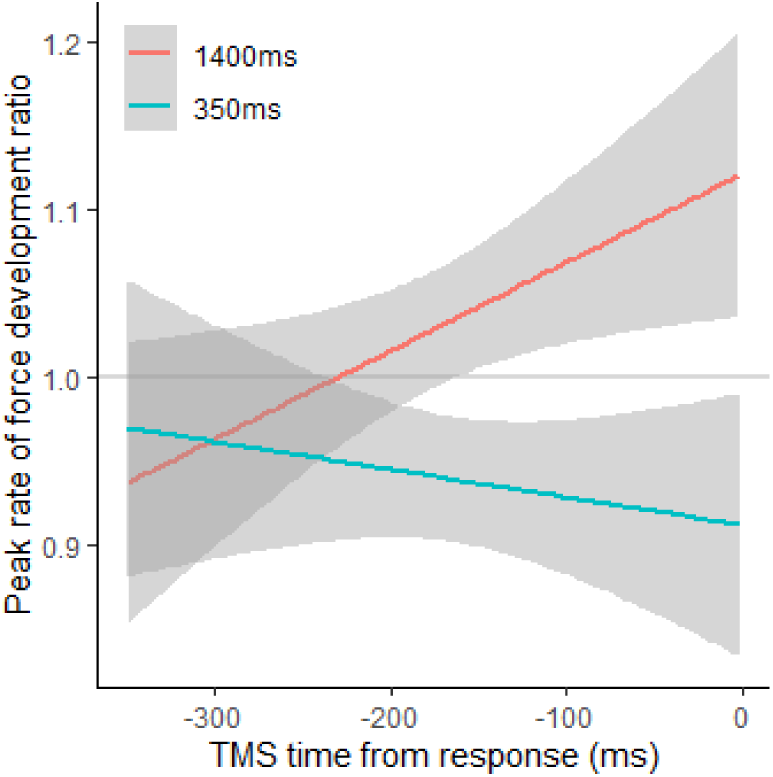
Peak rate of force development ratios as a function of the time of TMS administration from movement onset, for each urgency condition.

**FIGURE 6.**
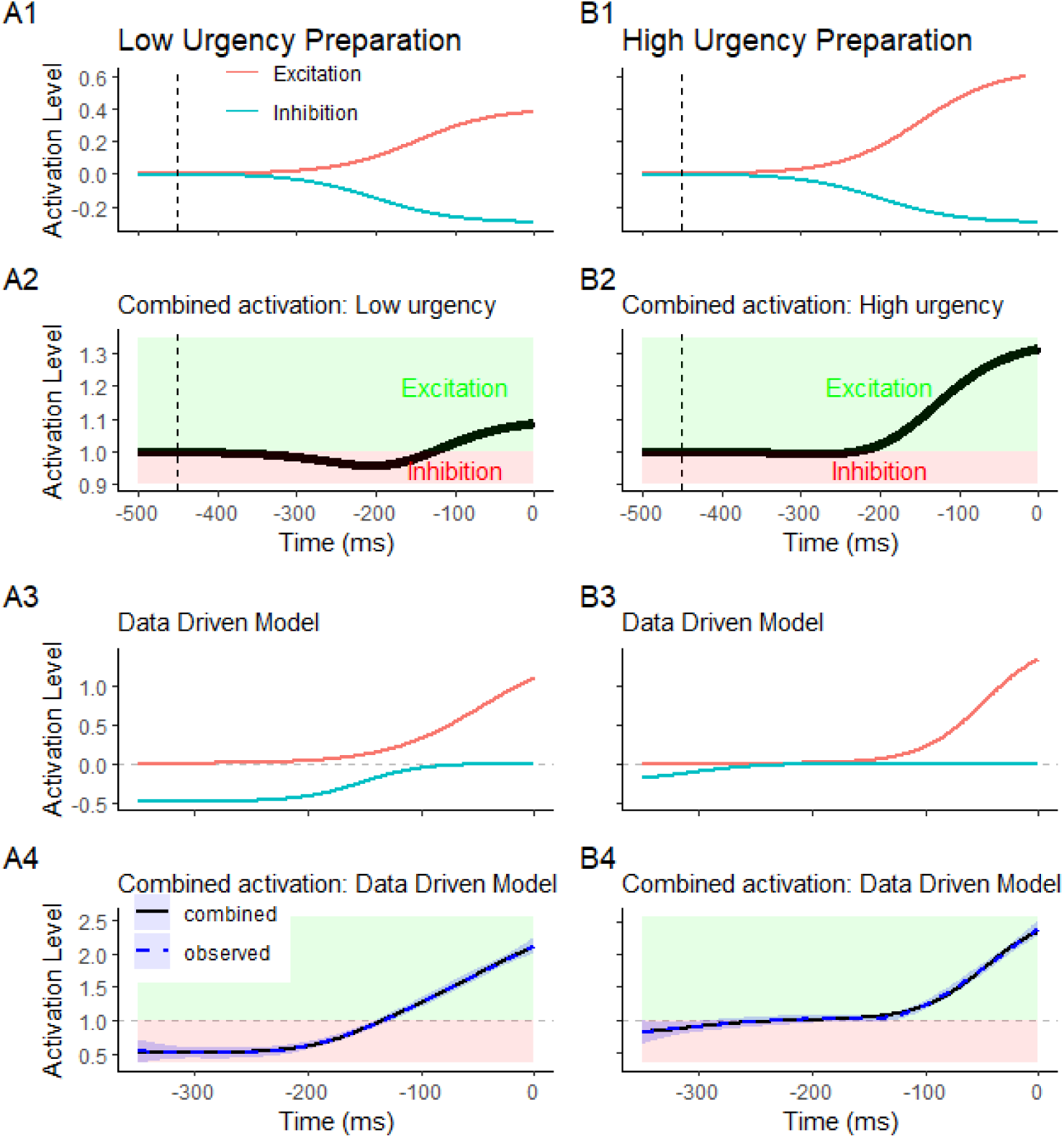
Modelled activation levels during movement preparation for low (A) and high (B) urgency conditions. A1-B1 Show inhibitory and excitatory processes for our conceptual model for which urgency modulates the growth rate of the excitatory sigmoid function. A2-B2 shows the combined net activation considering both inhibitory and excitatory inputs under low and high urgency. The blue line represents inhibitory processes and red line represents excitatory processes. Vertical dashed lines indicate when these processes are starting. The green shaded line represents net excitation and the red shaded lines represents net inhibition. A3-B3 show inhibitory and excitatory sigmoid functions with parameters fine-tuned via Maximum Likelihood Estimation, for which inhibition is constrained by urgency. These processes additively produce the combined net activation (A4-B4), which is consistent with observed MEPs.

## 4.0 Discussion

Suppression of CS excitability prior to action initiation has been repeatedly demonstrated in typical studies of movement preparation (Davranche et al., 2007; Duque et al., 2010; Duque & Ivry, 2009; Hannah et al., 2018; Hasbroucq et al., 1997, 1999; Ibáñez et al., 2020; Klein et al., 2016; Lebon et al., 2016; Marinovic et al., 2015; McInnes et al., 2021; Nguyen et al., 2021; Starr et al., 1988; Zaaroor et al., 2003). While it has been suggested that this process may be a necessary component of action preparation, recent data have suggested that the motor system can forego suppression during high-urgency contexts (McInnes et al., 2021). In the present study, we sought to both replicate these findings and extend them by examining how the temporal dynamics of CS inhibition might shift over the course of preparation in high-urgency contexts. We confirmed that preparatory suppression is absent under high urgency conditions, with no signs of a temporal shift of preparation processes under this context. We also observed evidence to support the notion that preparatory suppression can protect motor performance from interference.

In more detail, we modified the duration of time available to prepare an anticipatory action and applied single-pulse TMS at two times relative to the expected time of movement onset. The data here replicate the previously observed context-dependent modulation of CS excitability: TMS applied 250 ms prior to the time of expected movement onset indicated a suppression of CS excitability when preparing a low-urgency action, but not when preparing a high-urgency one. To investigate the possibility of a temporal shift in CS inhibition during high-urgency preparation, we applied TMS at 150 ms prior to the expected time of movement onset. Under low-urgency preparation, the resulting MEPs were no different from baseline, suggesting that the CS tract undergoes a disinhibition at this point of the preparatory process. However, TMS delivered at -150 ms before movement onset in the high urgency condition led to a facilitation of MEPs relative to baseline. Therefore, these data provide no evidence for a temporal shift in the time course of preparatory inhibition during high-urgency.

We further modelled the context-dependent time course of CS excitability by examining MEPs locked to the time of movement onset, rather than cue onset. Again, these data provided no evidence for a suppression of CS excitability when preparing a high-urgency action. In addition, the facilitation of CS excitability closer to action initiation was indicative of previously reported CS facilitation from approximately 100 ms prior to movement onset (Chen et al., 1998; Leocani et al., 2000; Starr et al., 1988; Zaaroor et al., 2003). Modelling of MEPs elicited over the course of low-urgency preparation and characterising the change in CS excitability as a function of time confirmed these findings. Up to approximately 238 ms prior to action onset, the net CS excitability was supressed during low-urgency preparation. Following this, CS excitability returned to baseline levels between 238 – 182 ms prior to movement onset, reflecting a change in the net balance between excitatory and inhibitory processes leading to movement execution. Finally, from 118 ms up until the time of movement onset, there was a significant net excitation of the CS tract. These data demonstrate the temporally tied modulation of CS excitability over the course of preparation in typical (low temporally constrained) laboratory experiments (Davranche et al., 2007, 2007; Duque et al., 2017; Duque & Ivry, 2009; Starr et al., 1988; Zaaroor et al., 2003).

We have previously demonstrated that motor execution is disrupted by both TMS and loud sounds when preparatory inhibition is absent, particularly under constrained preparation times (McInnes et al., 2021). Consistent with this, our cue-locked analysis with TMS showed a reduction in movement vigour when TMS was administered 250 ms prior to cue onset under high urgency preparation. This disruption was not observed in the low-urgency condition, where sufficient preparation time allowed for engagement in strategic premovement inhibition. Aligned with previous findings where acoustic stimuli increased response vigour (McInnes et al., 2021), our response-locked analysis of motor output suggested that TMS may facilitate actions when administered closer to the time of movement onset for low-urgency actions, and lead to a reduction of vigour under a high-urgency to move.

### 4.1 A conceptual model to understand the net activation of the corticospinal system under different urgency scenarios

Here we consider the extent to which the results of our study are consistent with a simple conceptual model of the temporal dynamics of corticospinal excitability during movement preparation under high and low urgency conditions. The model integrates excitatory and inhibitory processes, which we propose are related to glutamatergic and GABAergic inputs, respectively. For simplicity, we initially imagined a conceptual model where 1) inhibitory and excitatory processes are both automatically triggered by a go signal, and 2) the dynamics of both processes resemble a sigmoid function (Teka et al., 2017). Thus, the activation of inhibitory and excitatory processes can each be described by the sigmoid function:

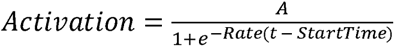

where *A* represents the magnitude of activation, *Rate* denotes the rate of growth in the sigmoid curve, *t* is time relative to movement onset, and *StartTime* denotes a shift of the sigmoid curve across time. The resulting inhibitory activation *Activation_I_* and excitatory activation *Activation_E_* are additive with baseline activation *Activation_B_* (a random variable with a mean = 1) to produce the net combined activation, *Activation_Net_*, which can be measured in MEPs:

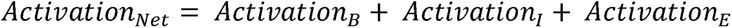

It is clear that there are a multitude of functions and dynamics that could be combined to explain the lack of preparatory suppression under high urgency to move and these will be discussed subsequently. In this conceptual model shown in **FIGURE 6**, the inhibitory process starts to develop at a faster rate at an earlier time than the excitatory process but reaches a lower amplitude overall. Under low temporal urgency to move (**FIGURE 6A1-A2**), this model would result in a net activation (Excitation + Inhibition) in which the net balance is inhibitory (greater inhibition) but reverts to excitatory as the processes unfold. This would explain why under low urgency to move, we and others have observed MEP suppression around 200 ms to movement onset (Davranche et al., 2007; Duque et al., 2010; Duque & Ivry, 2009; Hannah et al., 2018; Hasbroucq et al., 1997, 1999; Ibáñez et al., 2020; Klein et al., 2016; Lebon et al., 2016; Marinovic et al., 2015; McInnes et al., 2021; Starr et al., 1988; Zaaroor et al., 2003). Of more relevance here is what causes a lack of preparatory suppression of the corticospinal system under high urgency to move. In this scenario, we considered that the dynamics of inhibitory and excitatory processes are maintained. However, excitatory inputs are sensitive to the urgency to move. In **FIGURE 6B1-B2**, we simply multiplied the rate of excitation by an arbitrary factor (1.6 in the example shown in **FIGURE 6B1**), which was sufficient to completely eliminate any net inhibitory effect until the time of movement onset. The sensitivity of the excitatory process to urgency is well aligned with the results obtained by Thura and Cisek (2016) in behaving monkeys, where they found that primary motor cortex (M1) neurons displayed higher baseline firing rates and stronger modulation by accumulating evidence in trials where speed was emphasised over accuracy. Of course, it should be noted that although urgency only affects excitatory inputs in our model, it is possible that there might be a simultaneous decrease in inhibitory inputs to the spinal cord —our model only reflects the net activity and its peak is much higher under high than low urgency scenarios. However, this qualitative feature of the model would suggest that increased excitatory input related to the prepared response could lead to more forceful responses. In this regard, experiments have reported increased response vigour under high urgency to move (Mattes & Ulrich, 1997; McInnes et al., 2021; Thura & Cisek, 2016).

The conceptual model we propose assumes that inhibitory and excitatory processes are automatically engaged by a go-signal whenever an action needs to be initiated. An alternative could be that inhibitory and excitatory processes might be triggered by different signals. For example, when preparation time is sufficient, inhibitory processes are engaged early in preparation, perhaps before the excitatory process as a part of generic temporal preparation for action 27/12/2023 10:47:00 AM. However, when preparation times are short, the delayed engagement of inhibitory processes may either not occur, or start too late relative to the excitatory process to effectively reduce corticospinal excitability. In this case, the preparatory processes are relatively independent, potentially reflecting the processes of movement initiation and specification as put forward by Haith and colleagues (2016) (see also Niemi & Näätänen, 1981; Nobre et al., 2007). Under this independence assumption, the net activation at any timepoint is the same as for the conceptual model shown in **FIGURE 6B2**, the net activity between excitation and inhibition. This simple alternative conceptual model would also capture the pattern of MEP results observed in both our current study and previous work (McInnes et al., 2021; Ibáñez et al., 2020).

To establish the utility of this model, we examined the fit of the combined net activation, considering the sigmoid inhibitory and excitatory processes, to observed net activation. To this end, we used Maximum Likelihood Estimation (MLE) to fine-tune the parameter values of the inhibitory and excitatory sigmoid functions. We optimised parameters to find the net activation which maximises the negative log likelihood of observing the pattern of MEPs we report here. This data-driven model with optimised parameters supports an asynchronous inhibitory/excitatory process, where the inhibitory process is engaged before the excitatory process (**FIGURE 6A3**). Therefore, inhibitory processes may not be engaged when preparation time is constrained (**FIGURE 6B3**). Supporting our approach of modelling activation as a sigmoid function, the resulting net activation of these processes (**FIGURE 6A4, 6B4**) provide a strong fit with observed MEPs. However, given that we measured MEPs to single pulse TMS, which provides only a metric of net activation, it is important to note that we cannot definitively rule out that the combined activation is a product of latent opposing inhibitory and excitatory processes that start simultaneously.

We have previously proposed that preparatory suppression serves to protect upcoming responses from interference from internal or external events that could disrupt them. We have demonstrated both here and in previous work that responses prepared under high-urgency scenarios are less likely to be disrupted, and might actually be facilitated, by external stimuli. However, if this process is important to guarantee proper movement execution, why does it take relatively long to happen? According to the first model discussed above, where the inhibitory and excitatory processes are triggered by a common go-signal. The lack of protective suppression under high urgency conditions is simply an unwanted consequence that reflects the limitations of the motor system. In contrast, under the assumption that these processes are triggered by different signals, one could suggest that preparatory protection is relatively slow because it depends on attentional processes driven by the prefrontal cortex (PFC). This may be mediated by GABAergic inhibition (particularly, GABA_B;_ Ghosal et al., 2017; Kihara et al., 2016; Schmidt-Wilcke et al., 2018) and could be linked to automatic and attention-driven processes observed in other domains (e.g., social cognition) operating in tandem but being modulated by different factors as seen in disorders like schizophrenia (Langdon et al., 2017).

What is the evidence that attention may impact motor related neural processes preceding movement onset? Rowe et al. (2002) showed that attention to actions increased activity not only in the PFC but also in the premotor and parietal cortices. They also showed that the effective connectivity between the dorsal PFC and the M1 was enhanced when attention was directed to actions. These results indicate that the PFC can serve as a hub for cognitive control that influences motor processes via attentional modulation. This lends some validity to a conceptual model where inhibition and excitation can be triggered by independent signals and subject to attentional modulation orchestrated by the PFC. Do we have any evidence that conditions affecting PFC function can interfere with preparatory suppression? In conditions like Parkinson’s disease (PD), which is associated with cognitive dysfunction and can affect patients early on (Dirnberger & Jahanshahi, 2013), there is some evidence supporting this idea. More relevant to our conceptual model, recent work by Wilhelm et al. (2022) found that PD patients display reduced CS suppression during movement preparation in comparison to healthy control individuals. Relatedly, alcohol dependent individuals and binge drinkers, who commonly show abnormalities in prefrontal networks, show a similar impairment of CS suppression during action preparation (Grandjean & Duque, 2020; Quoilin et al., 2018). Further research on preparatory suppression under different temporal constraints will be important to advance our knowledge of this phenomenon, with potential implications to movement rehabilitation.

## 5.0 Conclusion

In conclusion, we have demonstrated that actions prepared under high-urgency undergo a different modulation of CS excitability from those that are prepared under low-urgency conditions. Specifically, under high temporal constraints, when there is little time available to prepare an action, the CS tract does not undergo the suppression that is typically observed under low temporal constraints. We considered two physiologically plausible yet simple conceptual models that can explain the dynamic transitions from suppression to facilitation of the corticospinal pathway. As real-world actions often allow little time for preparation, the modulation of CS pathways and motor output by urgency may provide a more translational paradigm to investigate functional motor control.

## 6.0 Funding

This work was funded by the Australian Research Council, DP180100394 to W.M, and DE190100694 to AMV.

## Notes

### Competing Interest Statement

The authors have declared no competing interest.

## References

Chen, R., Yaseen, Z., Cohen, L. G., & Hallett, M. (1998). Time course of corticospinal excitability in reaction time and self-paced movements. Annals of Neurology, 44(3), 317–325. 10.1002/ana.410440306

Davranche, K., Tandonnet, C., Burle, B., Meynier, C., Vidal, F., & Hasbroucq, T. (2007). The dual nature of time preparation: Neural activation and suppression revealed by transcranial magnetic stimulation of the motor cortex. European Journal of Neuroscience, 25(12), 3766–3774. 10.1111/j.1460-9568.2007.05588.x

Dirnberger, G., & Jahanshahi, M. (2013). Executive dysfunction in Parkinson’s disease: A review. Journal of Neuropsychology, 7(2), 193–224. 10.1111/jnp.12028

Duque, J., Greenhouse, I., Labruna, L., & Ivry, R. B. (2017). Physiological markers of motor inhibition during human behavior. Trends in Neurosciences, 40(4), 219–236. 10.1016/j.tins.2017.02.006

Duque, J., & Ivry, R. B. (2009). Role of corticospinal suppression during motor preparation. Cerebral Cortex, 19(9), 2013–2024. 10.1093/cercor/bhn230

Duque, J., Lew, D., Mazzocchio, R., Olivier, E., & Ivry, R. B. (2010). Evidence for two concurrent inhibitory mechanisms during response preparation. Journal of Neuroscience, 30(10), 3793–3802. 10.1523/JNEUROSCI.5722-09.2010

Ghosal, S., Hare, B., & Duman, R. S. (2017). Prefrontal Cortex GABAergic Deficits and Circuit Dysfunction in the Pathophysiology and Treatment of Chronic Stress and Depression. Current Opinion in Behavioral Sciences, 14, 1–8. 10.1016/j.cobeha.2016.09.012

Grandjean, J., & Duque, J. (2020). A TMS study of preparatory suppression in binge drinkers. NeuroImage: Clinical, 28, 102383. 10.1016/j.nicl.2020.102383

Haith, A. M., Pakpoor, J., & Krakauer, J. W. (2016). Independence of movement preparation and movement initiation. Journal of Neuroscience, 36(10), 3007–3015. 10.1523/JNEUROSCI.3245-15.2016

Hannah, R., Cavanagh, S. E., Tremblay, S., Simeoni, S., & Rothwell, J. C. (2018). Selective suppression of local interneuron circuits in human motor cortex contributes to movement preparation. Journal of Neuroscience, 38(5), 1264–1276. 10.1523/JNEUROSCI.2869-17.2017

Hasbroucq, T., Kaneko, H., Akamatsu, M., & Possamaï, C. A. (1997). Preparatory inhibition of cortico-spinal excitability: A transcranial magnetic stimulation study in man. Cognitive Brain Research, 5(3), 185–192. 10.1016/S0926-6410(96)00069-9

Hasbroucq, T., Kaneko, H., Akamatsu, M., & Possamaï, C. A. (1999). The time-course of preparatory spinal and cortico-spinal inhibition: An H-reflex and transcranial magnetic stimulation study in man. Experimental Brain Research, 124(1), 33–41. 10.1007/s002210050597

Ibáñez, J., Hannah, R., Rocchi, L., & Rothwell, J. C. (2020). Premovement suppression of corticospinal excitability may be a necessary part of movement preparation. Cerebral Cortex, 30(5), 2910–2923. 10.1093/cercor/bhz283

Kihara, K., Kondo, H. M., & Kawahara, J. I. (2016). Differential Contributions of GABA Concentration in Frontal and Parietal Regions to Individual Differences in Attentional Blink. Journal of Neuroscience, 36(34), 8895–8901. 10.1523/JNEUROSCI.0764-16.2016

Klein, P.-A., Duque, J., Labruna, L., & Ivry, R. B. (2016). Comparison of the two cerebral hemispheres in inhibitory processes operative during movement preparation. NeuroImage, 125, 220–232. 10.1016/j.neuroimage.2015.10.007

Kleiner, M., Brainard, D., Pelli, D., Ingling, A., Murray, R., & Broussard, C. (2007). What’s new in psychtoolbox-3. Perception, 36(14), 1–16.

Langdon, R., Seymour, K., Williams, T., & Ward, P. B. (2017). Automatic attentional orienting to other people’s gaze in schizophrenia. Quarterly Journal of Experimental Psychology, 70(8), 1549–1558. 10.1080/17470218.2016.1192658

Lebon, F., Greenhouse, I., Labruna, L., Vanderschelden, B., Papaxanthis, C., & Ivry, R. B. (2016). Influence of Delay Period Duration on Inhibitory Processes for Response Preparation. Cerebral Cortex, 26(6), 2461–2470. 10.1093/cercor/bhv069

Leocani, L., Cohen, L. G., Wassermann, E. M., Ikoma, K., & Hallett, M. (2000). Human corticospinal excitability evaluated with transcranial magnetic stimulation during different reaction time paradigms. Brain, 123(6), 1161–1173. 10.1093/brain/123.6.1161

Marinovic, W., Flannery, V., & Riek, S. (2015). The effects of preparation and acoustic stimulation on contralateral and ipsilateral corticospinal excitability. Human Movement Science, 42, 81–88. 10.1016/j.humov.2015.05.003

Mattes, S., & Ulrich, R. (1997). Response force is sensitive to the temporal uncertainty of response stimuli. Perception and Psychophysics, 59(7), 1089–1097. 10.3758/BF03205523

McInnes, A. N., Lipp, O. V., Tresilian, J. R., Vallence, A. M., & Marinovic, W. (2021). Premovement inhibition can protect motor actions from interference by response-irrelevant sensory stimulation. Journal of Physiology, 599(18), 4389–4406. 10.1113/JP281849

Nguyen, A. T., Jacobs, L. A., Tresilian, J. R., Lipp, O. V., & Marinovic, W. (2021). Preparatory suppression and facilitation of voluntary and involuntary responses to loud acoustic stimuli in an anticipatory timing task. Psychophysiology, 58(2), e13730. 10.1111/psyp.13730

Niemi, P., & Näätänen, R. (1981). Foreperiod and simple reaction time. Psychological Bulletin, 89(1), 133–162. 10.1037/0033-2909.89.1.133

Nobre, A., Correa, A., & Coull, J. (2007). The hazards of time. Current Opinion in Neurobiology, 17(4), 465–470. 10.1016/j.conb.2007.07.006

Quoilin, C., Wilhelm, E., Maurage, P., de Timary, P., & Duque, J. (2018). Deficient inhibition in alcohol-dependence: Let’s consider the role of the motor system! Neuropsychopharmacology, 43(9), Article 9. 10.1038/s41386-018-0074-0

Rossi, S., Antal, A., Bestmann, S., Bikson, M., Brewer, C., Brockmöller, J., Carpenter, L. L., Cincotta, M., Chen, R., Daskalakis, J. D., Di Lazzaro, V., Fox, M. D., George, M. S., Gilbert, D., Kimiskidis, V. K., Koch, G., Ilmoniemi, R. J., Pascal Lefaucheur, J., Leocani, L., … Hallett, M. (2021). Safety and recommendations for TMS use in healthy subjects and patient populations, with updates on training, ethical and regulatory issues: Expert Guidelines. Clinical Neurophysiology, 132(1), 269–306. 10.1016/J.CLINPH.2020.10.003

Rossini, P. M., Barker, A. T., Berardelli, A., Caramia, M. D., Caruso, G., Cracco, R. Q., Dimitrijević, M. R., Hallett, M., Katayama, Y., Lücking, C. H., Maertens de Noordhout, A. L., Marsden, C. D., Murray, N. M. F., Rothwell, J. C., Swash, M., & Tomberg, C. (1994). Non-invasive electrical and magnetic stimulation of the brain, spinal cord and roots: Basic principles and procedures for routine clinical application. Report of an IFCN committee. Electroencephalography and Clinical Neurophysiology, 91(2), 79–92. 10.1016/0013-4694(94)90029-9

Rowe, J., Friston, K., Frackowiak, R., & Passingham, R. (2002). Attention to action: Specific modulation of corticocortical interactions in humans. NeuroImage, 17(2), 988–998.

Schmidt-Wilcke, T., Fuchs, E., Funke, K., Vlachos, A., Müller-Dahlhaus, F., Puts, N. A. J., Harris, R. E., & Edden, R. A. E. (2018). GABA—from Inhibition to Cognition: Emerging Concepts. The Neuroscientist, 24(5), 501–515. 10.1177/1073858417734530

Starr, A., Caramia, M., Zarola, F., & Rossini, P. M. (1988). Enhancement of motor cortical excitability in humans by non-invasive electrical stimulation appears prior to voluntary movement. Electroencephalography and Clinical Neurophysiology, 70(1), 26–32. 10.1016/0013-4694(88)90191-5

Teka, W. W., Hamade, K. C., Barnett, W. H., Kim, T., Markin, S. N., Rybak, I. A., & Molkov, Y. I. (2017). From the motor cortex to the movement and back again. PLoS ONE, 12(6), e0179288. 10.1371/journal.pone.0179288

Thura, D., & Cisek, P. (2016). Modulation of Premotor and Primary Motor Cortical Activity during Volitional Adjustments of Speed-Accuracy Trade-Offs. The Journal of Neuroscience, 36(3), 938–956. 10.1523/JNEUROSCI.2230-15.2016

Wilhelm, E., Quoilin, C., Derosiere, G., Paço, S., Jeanjean, A., & Duque, J. (2022). Corticospinal Suppression Underlying Intact Movement Preparation Fades in Parkinson’s Disease. Movement Disorders: Official Journal of the Movement Disorder Society, 37(12), 2396–2406. 10.1002/mds.29214

Zaaroor, M., Pratt, H., & Starr, A. (2003). Time course of motor excitability before and after a task-related movement. Neurophysiologie Clinique, 33, 130–137. 10.1016/S0987-7053(03)00029-7

